# Activity in the Fronto-Parietal Multiple-Demand Network is Robustly Associated with Individual Differences in Working Memory and Fluid Intelligence

**DOI:** 10.1101/110270

**Authors:** Moataz Assem, Idan Asher Blank, Zachary Mineroff, Ahmet Ademoglu, Evelina Fedorenko

**Author notes:** corresponding authors (MA) and (EF).

## Abstract

Numerous brain lesion and fMRI studies have linked individual differences in executive abilities and fluid intelligence to brain regions of the fronto-parietal “multiple-demand” (MD) network. Yet, fMRI studies have yielded conflicting evidence as to whether better executive abilities are associated with stronger or weaker MD activations and whether this relationship is restricted to the MD network. Here, in a large-sample (n=216) fMRI investigation, we found that stronger activity in MD regions – functionally defined in individual participants – was robustly associated with more accurate and faster responses on a spatial working memory task performed in the scanner, as well as fluid intelligence measured independently (n=114). In line with some prior claims about a relationship between language and fluid intelligence, we also found a weak association between activity in the brain regions of the left fronto-temporal language network during an independent passive reading task, and performance on the working memory task. However, controlling for the level of MD activity abolished this relationship, whereas the MD activity-behavior association remained highly reliable after controlling for the level of activity in the language network. Finally, we demonstrate how unreliable MD activity measures, coupled with small sample sizes, could falsely lead to the opposite, negative, association that has been reported in some prior studies. Taken together, these results demonstrate that a core component of individual differences variance in executive abilities and fluid intelligence is selectively and robustly positively associated with the level of activity in the MD network, a result that aligns well with lesion studies.

## Introduction

General cognitive abilities, such as fluid intelligence, and the tightly linked executive abilities, are among the best predictors of academic achievement and professional success (Gottfredson, 2002; Kuncel & Hezlett, 2010; Plomin & Deary, 2015). These abilities are thought to rely on a network of bilateral frontal and parietal brain regions. Selective damage to these regions is associated with disorganized executive behavior and significant loss of fluid intelligence (Duncan, Burgess, & Emslie, 1995; Glascher et al., 2010; Roca et al., 2010; Warren et al., 2014; Woolgar, Duncan, Manes, & Fedorenko, 2018; Woolgar et al., 2010). Similar frontal and parietal regions are activated in brain imaging studies during diverse demanding tasks, including manipulations of working memory, fluid reasoning, selective attention, set shifting, response inhibition, and novel problem solving inter alia (Assem, Glasser, Essen, & Duncan, 2020; Michael W. Cole & Schneider, 2007; Dosenbach et al., 2006; Duncan, 2000, 2010; Duncan & Owen, 2000; Fedorenko, Duncan, & Kanwisher, 2013; Geake & Hansen, 2005; Vakhtin, Ryman, Flores, & Jung, 2014). We refer to this set of brain regions as the “multiple-demand” (MD) network (following Duncan, 2013, 2010) given their sensitivity to multiple task demands. The MD network includes lateral and dorsomedial frontal areas, anterior insular areas, and areas along the intra-parietal sulcus (Assem et al., 2020; Fedorenko et al., 2013), and these areas form a functionally integrated system as evidenced by strong synchronization during naturalistic cognition (Assem et al., 2020; Blank, Kanwisher, & Fedorenko, 2014; Paunov, Blank, & Fedorenko, 2019).

Prior fMRI studies have linked activity in the MD network with individual differences in executive abilities and fluid intelligence, but have left open the nature of this relationship. First, conflicting claims have been made regarding the direction of brain-behavior correlations across individuals. On the one hand, some have found that stronger MD activation is associated with worse performance on executive tasks and lower IQ (Basten, Hilger, & Fiebach, 2015; Deary, Penke, & Johnson, 2010; Dunst et al., 2014; Haier et al., 1988; Neubauer & Fink, 2009; Rypma et al., 2006; Rypma & Esposito, 2000; Santarnecchi, Galli, Polizzotto, Rossi, & Rossi, 2014; Stern, Gazes, Razlighi, Steffener, & Habeck, 2018). Such studies have typically advocated a “neural efficiency” explanation: smarter individuals can use fewer neural resources to achieve the same level of performance. On the other hand, others have found the opposite pattern, where stronger MD activation is associated with better executive task performance and higher IQ (Basten, Stelzel, & Fiebach, 2013; Burgess, Gray, Conway, & Braver, 2011; Choi et al., 2008; M. W. Cole, Yarkoni, Repovs, Anticevic, & Braver, 2012; Gray, Chabris, & Braver, 2003; Lee et al., 2006; Tschentscher, Mitchell, & Duncan, 2017). In an attempt to reconcile these conflicting findings, some have suggested that the direction of the correlation may depend on task difficulty with “neural efficiency” (i.e., a negative association between MD activity and performance) observed in easier tasks, and positive associations observed during more complex tasks (Neubauer & Fink, 2009; Sripada, Angstadt, Rutherford, Taxali, & Shedden, 2020).

Relatedly, superior executive abilities and higher IQ have been reported to correlate with stronger synchronization (typically, estimated during rest e.g. Fox et al., 2005) among the MD brain regions (M. W. Cole et al., 2012; Dubois, Galdi, Paul, & Adolphs, 2018; Ferguson, Anderson, & Spreng, 2017; Finn et al., 2015; Hearne, Mattingley, & Cocchi, 2016; Smith et al., 2015), although most of these studies have relied on the same resting-state Human Connectome Project (HCP) dataset (Smith et al., 2013). Fewer studies have reported weaker synchronization in such individuals (Santarnecchi et al., 2014; van den Heuvel, Stam, Kahn, & Hulshoff Pol, 2009).

A second open question concerns the specificity of this relationship to the MD network. Challenging the idea that executive functions are selectively tied to this network, a number of fMRI studies have also linked individual differences in executive abilities and fluid intelligence with activity in other brain regions/networks, including occipito-temporal areas ((Haier, White, & Alkire, 2003; Park, Carp, Hebrank, Park, & Polk, 2010) but see (Assem et al., 2020; Sani, McPherson, Stemmann, Pestilli, & Freiwald, 2019) for evidence that these regions may belong to an extended MD network), the default mode network (DMN) (Lipp et al., 2012; Smith et al., 2015), or the degree of MD-DMN differentiation (Sripada et al., 2020). A recent study used 7 fMRI tasks from the HCP dataset to demonstrate that task activation levels in many brain regions can, to some extent, predict individual differences in general intelligence, though critically, MD regions—engaged by executive function tasks—are the best predictors (Sripada et al., 2020). In contrast, another recent study using the HCP resting-state dataset showed that the strength of inter-region correlations in most brain networks predicts general intelligence, and to a similar extent (Dubois et al., 2018). A key potential limitation of these studies is that, like the above-mentioned studies, they rely exclusively on the HCP dataset and are yet to be replicated in independent data.

These apparently discrepant results could reflect the complexity of the brain-behavior relationship in the domain of executive abilities, with perhaps multiple underlying cognitive constructs (Miyake & Friedman, 2012) and neural mechanisms contributing to their implementation. However, a number of methodological limitations plague previous studies and may, instead, explain away some of these discrepancies. *First*, many earlier studies have used small numbers of participants (as low as n=8) and/or transformed continuous behavioral measures into categorical variables (e.g., high- vs. low-performing participants). Both of these factors can produce inflated or spurious relationships (Haier et al., 1988; Lee et al., 2006; Rypma et al., 2006; Rypma & Esposito, 2000; Wager et al., 2005). *Second*, most studies have failed to assess the reliability of the relevant behavioral and/or brain measures (e.g., the strength of the BOLD response, or the strength of inter-regional synchronization) – a critical prerequisite for relating behavioral and brain individual variability (Dubois & Adolphs, 2016). Both behavioral and brain measures have to be stable within individuals over time (e.g., across multiple runs of a task, or across tasks) (Mahowald & Fedorenko, 2016). This is especially important for studies using BOLD estimates based on contrasts of task relative to fixation, or resting-state inter-region synchronization measures, which may fail to isolate MD activity from general state variables, like motivation, arousal, or caffeine intake (Basten et al., 2013; M. W. Cole et al., 2012; Dubois et al., 2018; Dunst et al., 2014; Gray et al., 2003; Rypma et al., 2006; Rypma & Esposito, 2000; Smith et al., 2015; Stern et al., 2018; Wager et al., 2005). *Third*, almost all previously mentioned studies have failed to take into consideration individual variability in the precise locations of the MD regions (see (Assem et al., 2020; Blank, 2020; Fedorenko et al., 2013; Shashidhara, Spronkers, & Erez, 2020) for evidence of such variability). This variability leads to losses in sensitivity and functional resolution (Brett, Johnsrude, & Owen, 2002; Nieto-Castañón & Fedorenko, 2012; Saxe, Brett, & Kanwisher, 2006), and it also affects the interpretation of inter-regional functional synchronization findings (Bijsterbosch, Beckmann, Woolrich, Smith, & Harrison, 2019; Bijsterbosch et al., 2018). This problem is compounded by the proximity of MD areas to functionally distinct areas such as language-selective regions (Fedorenko, Duncan, & Kanwisher, 2012), which show no response to any demanding task other than language processing (Fedorenko, Behr, & Kanwisher, 2011; Fedorenko & Blank, 2020; Fedorenko & Varley, 2016; Monti, Parsons, & Osherson, 2012). And *fourth*, many studies have failed to adequately assess the selectivity of the relationship between MD activity and behavior (Choi et al., 2008; M. W. Cole et al., 2012; Dubois & Adolphs, 2016; Gray et al., 2003; Rypma et al., 2006). This is important given that trait variables (e.g., brain vascularization) are known to affect neural responses (e.g., Ainslie and Duffin, 2009; Kazan et al., 2016), so in order to argue that the MD network’s activity relates to individual differences in executive functions or fluid intelligence, it is important to demonstrate that activity in some other, control, brain region or network does not show a similar relationship.

To circumvent these limitations and rigorously test the relationship between MD activity and executive abilities and fluid intelligence, we conducted a large-scale fMRI study, where participants (n=216) performed a spatial working memory (WM) task that included a harder and an easier condition. We first established the reliability of the Hard>Easy (H>E) BOLD effect in the MD network (defined functionally in each participant individually (Fedorenko et al., 2013)), and then examined the relationship between the size of this effect and a) behavioral performance on the task (including in an independent run of data), and b) fluid intelligence (in a subset of participants, n=114). We further evaluated the selectivity of this MD-behavior relationship by examining fMRI responses in the left fronto-temporal language network while the same participants performed a language comprehension task (Fedorenko et al., 2010). This network serves as a good control because, on the one hand, the language network is robustly functionally distinct from the MD network (Blank et al., 2014; Diachek, Blank, Siegelman, Affourtit, & Fedorenko, 2020; Fedorenko & Blank, 2020; Mineroff, Blank, Mahowald, & Fedorenko, 2018), but on the other hand, language has long been implicated in abstract and flexible thought (e.g., Bickerton, 1995; Carruthers, 2002; Dennett, 1997; cf. Fedorenko and Varley, 2016), including some studies that have linked damage to the regions of this network to performance on some fluid reasoning tasks (e.g., Baldo et al., 2010; cf. Woolgar et al., 2018).

To foreshadow our results, we found that stronger (rather than weaker) MD responses were associated with better performance on the spatial WM task as well as higher fluid intelligence scores. The strength of activity in another large-scale network – the language network – did not explain any additional variability in WM task performance (i.e., it showed a weak correlation with behavior, which was eliminated once the level of MD activity was taken into account). Finally, we demonstrate how unreliable MD activity measures, coupled with small sample sizes, could lead to the opposite (negative) association between MD activity level and behavior as has been reported in the literature. These results align well with findings from lesion studies that have suggested that a substantial portion of the variance in executive abilities and fluid intelligence is strongly and selectively associated with frontal and parietal MD brain regions.

## Materials and Methods

### Participants

216 participants (age 23.6 ± 6.4, 136 males, 190 right handed, 13 left handed, 8 ambidextrous, 5 with missing handedness data) with normal or corrected-to-normal vision, students at Massachusetts Institute of Technology (MIT) and members of the surrounding community, participated for payment. All participants gave informed consent in accordance with the requirements of the Committee On the Use of Humans as Experimental Subjects (COUHES) at MIT.

### Experimental Paradigms

Participants performed a spatial working memory task in a blocked design (**Fig. 1**). Each trial lasted 8 seconds: within a 3×4 grid, a set of locations lit up in blue, one at a time for a total of 4 (easy condition) or two at a time for a total of 8 (hard condition). Participants were asked to keep track of the locations. At the end of each trial, they were shown two grids with some locations lit up and asked to choose the grid that showed the correct, previously shown locations by pressing one of two buttons. They received feedback on whether they answered correctly. Each participant performed two runs, with each run consisting of six 32-second easy condition blocks, six 32-second hard condition blocks, and four 16-second fixation blocks, for a total duration of 448s (7min 28s). Condition order was counterbalanced across runs.

**Figure 1.**
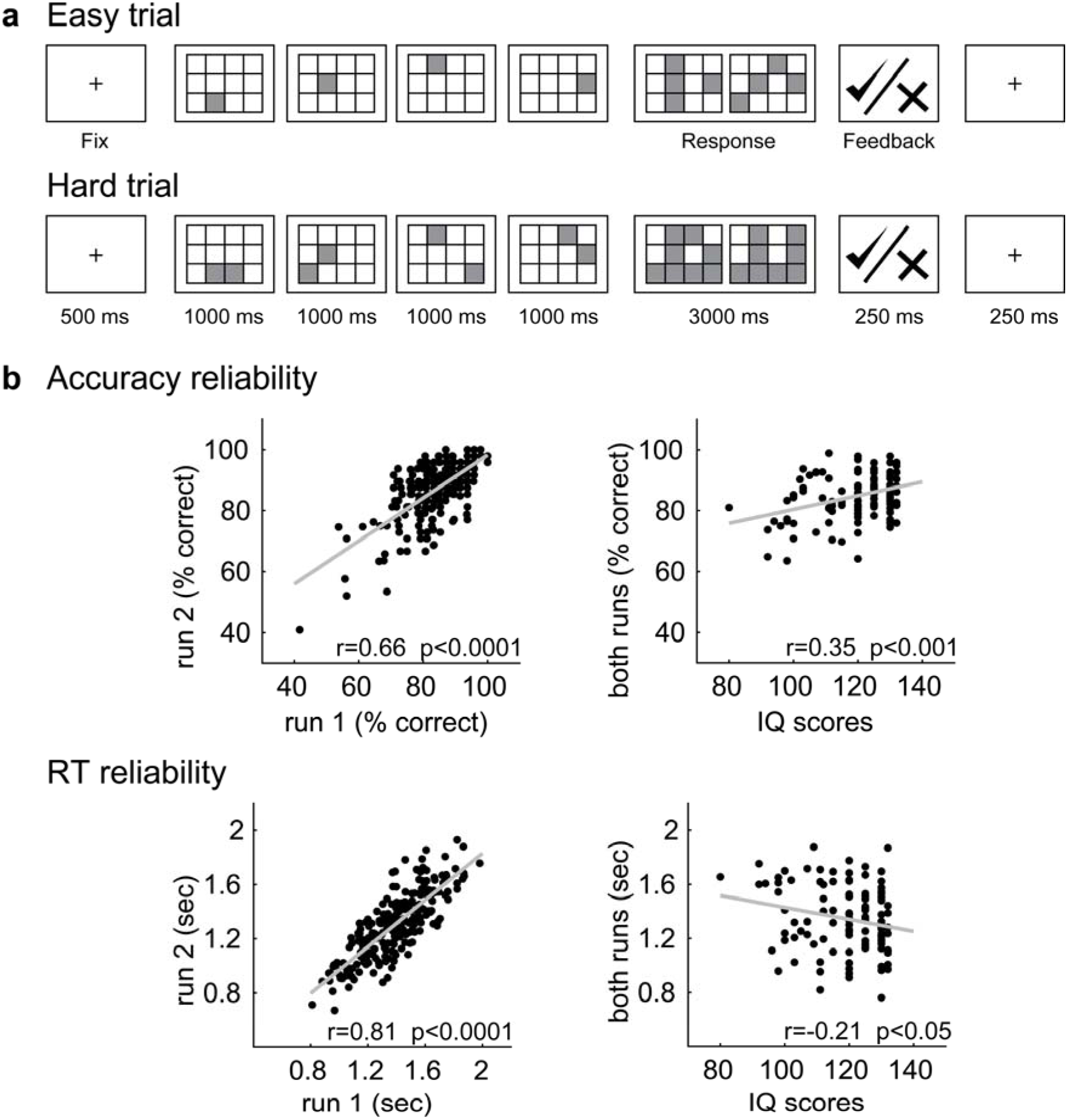
(a) Sample trials of the in-scanner spatial WM task, and (b) the reliability of its behavioral measures (averaging across the Easy and Hard conditions) across runs (in the full sample of n=216 participants) and with an independent measure of IQ (in a subset of n=114 participants).

In addition to the spatial working memory task, all participants performed a language localizer task (Fedorenko et al., 2010), used here to test the selectivity of the relationship between the MD network’s activity and behavior. The majority of the participants (n=182, 84.3%) passively read sentences and lists of pronounceable nonwords in a blocked design (see Table 1). The Sentences>Nonwords (S>N) contrast targets brain regions sensitive to high-level linguistic processing (Fedorenko et al., 2011, 2010). Each trial started with 100ms pre-trial fixation, followed by a 12-word-long sentence or a list of 12 nonwords presented on the screen one word/nonword at a time at the rate of 450ms per word/nonword. Then, a line drawing of a hand pressing a button appeared for 400ms, and participants were instructed to press a button whenever they saw the icon, and finally a blank screen was shown for 100ms, for a total trial duration of 6s. The button-press task was included to help participants stay alert and focused. Each block consisted of 3 trials and lasted 18s. Each participant performed two runs, with each run consisting of sixteen experimental blocks (eight per condition), and five fixation blocks (14s each), for a total duration of 358s (5min 58s). Condition order was counterbalanced across runs. The remaining 21 participants performed similar versions of the language localizer with minor differences in the timing and procedure, with one participant performing an auditory version of the localizer (see Table 1 for exact timings and procedures; we have previously established that the localizer contrast is robust to such differences (Fedorenko et al., 2010; Mahowald & Fedorenko, 2016; Scott, Gallée, & Fedorenko, 2017).

**Table 1.** Details of the design, materials, and procedure for the different variants of the language localizer task.

|  | Version |  |  |  |
| --- | --- | --- | --- | --- |
|  | A | B | C | D |
| Number of participants | 182 | 12 | 1 | 8 |
| Task (Passive Reading/Listening / Memory) | PR | M | PL | M |
| Words / nonwords per trial | 12 | 12 | variable | 12 |
| Trial duration (ms) | 6,000 | 6000 | 18000 | 6000 |
| Fixation | 100 | 300 | 0 | 300 |
| Presentation of each word / nonword | 450 | 350 | variable | 350 |
| Probe (M) + button press (M/PR) | 400 | 1000 | -- | 1000 |
| Fixation | 100 | 500 | 0 | 500 |
| Trials per block | 3 | 3 | 1 | 3 |
| Block duration (s) | 18 | 18 | 18 | 18 |
| Blocks per condition (per run) | 8 | 8 | 8 | 6 |
| Conditions | Sentences<br>Nonwords | Sentences<br>Nonwords | Intact<br>speech<br>Degraded<br>speech | Sentences<br>Nonwords<br>Word-lists<br>(not used<br>here) |
| Fixation block duration (s) | 14 | 18 | 14 | 18 |
| Number of fixation blocks per run | 5 | 5 | 5 | 4 |
| Total run time (s) | 358 | 378 | 358 | 396 |
| Number of runs | 2 | 2 | 2 | 2-3 |

Finally, most participants completed one or more additional experiments for unrelated studies. The entire scanning session lasted approximately 2 hours.

A subset of 114 participants performed the non-verbal component of KBIT (Kaufman & Kaufman, 2014) after the scanning session. The test consists of 46 items (of increasing difficulty) and includes both meaningful stimuli (people and objects) and abstract ones (designs and symbols). All items require understanding the relationships among the stimuli and have a multiple-choice format. If a participant answers 4 questions in a row incorrectly, the test is terminated, and the remaining items are marked as incorrect. The test is scored following the formal guidelines to calculate each participant’s IQ score.

### MRI data acquisition

Structural and functional data were collected on the whole-body 3 Tesla Siemens Trio scanner with a 32-channel head coil at the Athinoula A. Martinos Imaging Center at the McGovern Institute for Brain Research at MIT. T1-weighted structural images were collected in 128 axial slices with 1mm isotropic voxels (TR=2,530ms, TE=3.48ms). Functional, blood oxygenation level dependent (BOLD) data were acquired using an EPI sequence (with a 90° flip angle and using GRAPPA with an acceleration factor of 2), with the following acquisition parameters: thirty-one 4mm thick near-axial slices, acquired in an interleaved order with a 10% distance factor; 2.1mm x 2.1mm in-plane resolution; field of view of 200mm in the phase encoding anterior to posterior (A > P) direction; matrix size of 96mm x 96mm; TR of 2,000ms; and TE of 30ms. Prospective acquisition correction (Thesen, Heid, Mueller, & Schad, 2000) was used to adjust the positions of the gradients based on the participant’s motion one TR back. The first 10s of each run were excluded to allow for steady-state magnetization.

### FMRI data preprocessing and first-level analysis

FMRI data were analyzed using SPM5 and custom MATLAB scripts in volume space. (Note that first-level analyses have not changed much in later versions of SPM; we used an older version of the software here due to the use of these data in other projects spanning many years and hundreds of subjects; critical second-level analyses were performed using custom MATLAB scripts. We also verified using an independent dataset that estimates of neural activity extracted with SPM5- vs. SPM12-preprocessed and modeled data were extremely similar). Each subject’s data were motion corrected and then normalized into a common brain space (the Montreal Neurological Institute (MNI) template) and resampled into 2mm isotropic voxels. The data were then smoothed with a 4mm Gaussian filter (FWHM) and high-pass filtered (at 200s). The task effects in both the spatial WM task and in the language localizer task were estimated using a General Linear Model (GLM) in which each experimental condition was modeled with a separate boxcar regressor (with boxcars corresponding to blocks). For the working memory task, each run was modelled by one regressor for the easy blocks and one regressor for the hard blocks; similarly for the language task, each run was modelled by one regressor for sentence blocks and one regressor for non-word blocks. Regressors were convolved with the canonical hemodynamic response function (HRF). The model also included first-order temporal derivatives of these effects, as well as nuisance regressors representing entire experimental runs and offline-estimated motion parameters.

Fixation blocks in both tasks were not modeled and treated as the implicit baseline.

### MD fROIs definition and response estimation

To define the MD and language (see below) functional regions of interest (fROIs), we used the Group-constrained Subject-Specific (GSS) approach (Fedorenko et al., 2010). In particular, fROIs were constrained to fall within a set of “masks”, areas that corresponded to the expected gross locations of activation for the relevant contrast. For the MD fROIs, following Fedorenko et al. (Fedorenko et al., 2013) and Blank et al. (Blank et al., 2014), we used eighteen anatomical masks (Tzourio-Mazoyer et al., 2002) across the two hemispheres. These masks covered the portions of the frontal and parietal cortices where MD activity has been previously reported, including bilateral opercular inferior frontal gyrus (L/R IFGop), middle frontal gyrus (L/R MFG), orbital MFG (L/R MFGorb), insular cortex (L/R Insula), precentral gyrus (L/R PrecG), supplementary and presupplementary motor areas (L/R SMA), inferior parietal cortex (L/R ParInf), superior parietal cortex (L/R ParSup), and anterior cingulate cortex (L/R ACC) (**Fig. 2a**). It is worth noting, however, that a whole-brain GSS analysis (Fedorenko et al., 2010) performed on the Hard>Easy spatial WM activation maps of n=197 participants yields a set of functional masks that largely overlap with these anatomical parcels (e.g. Diachek et al., 2020). Within each mask, we selected the top 10% (as well as the top 20% and 30% for validation analyses, as described below) of most responsive voxels in each individual participant based on the *t-*values for the H>E spatial WM contrast. This top n% approach ensures that each fROI can be defined in every participant, and that the fROI sizes are identical across participants.

**Figure 2.**
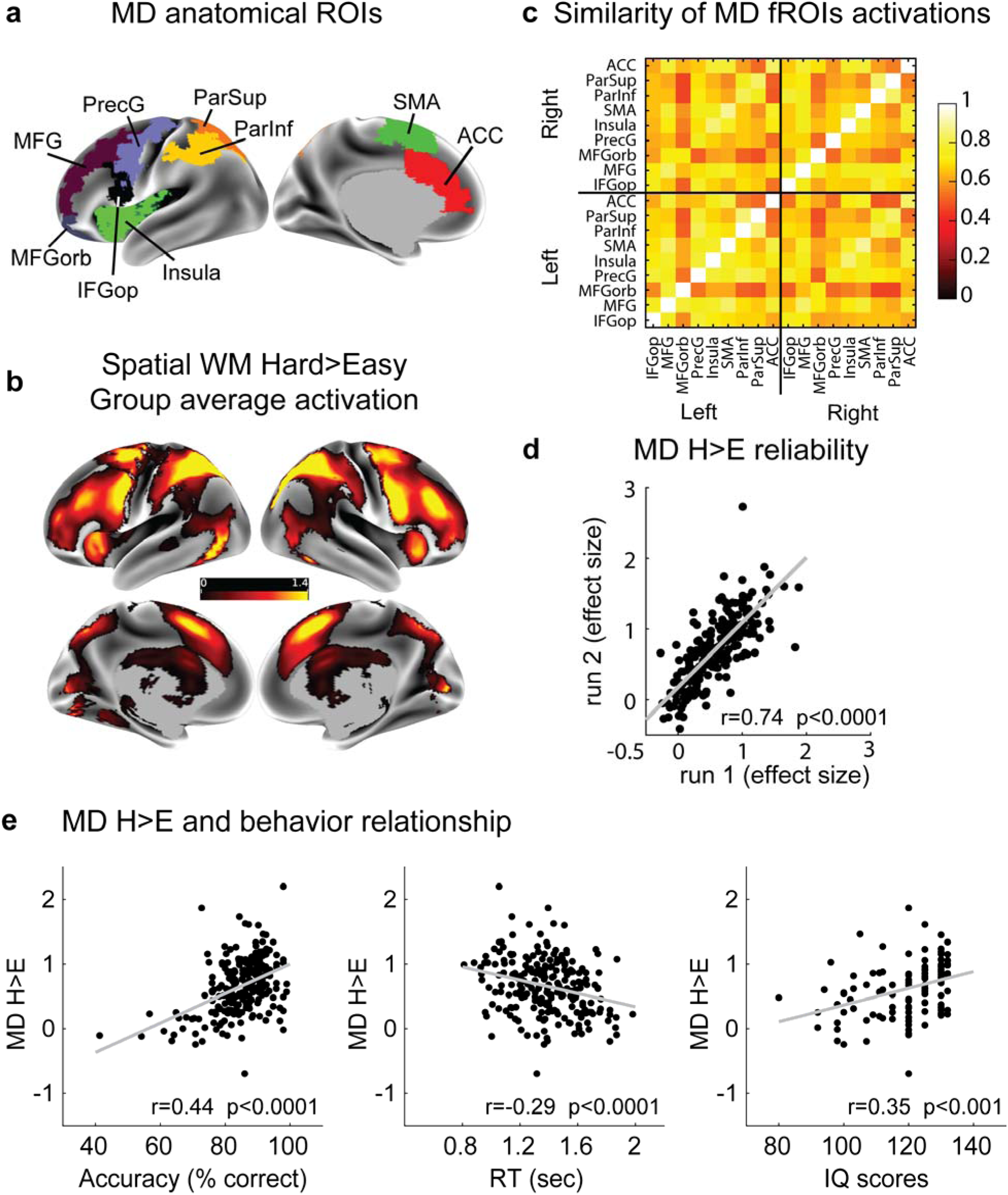
MD activity and behavior. **(a)** Surface projection of the volumetric anatomical masks used to constrain individual-specific functional activations. **(b)** Surface projection of the volumetric unthresholded group average activation map (beta estimates) for the spatial WM Hard>Easy (H>E) contrast. Please note that all analyses were performed in volume space, and surface projections—here and in other figures— are for illustrative purposes only and may include slight distortions resulting from volume-to-surface transformations. (Surface projection was performed using Connectome Workbench (humanconnectome.org/software/connectome-workbench) function “-volume-to-surface-mapping” using trilinear interpolation and a MNI reconstructed mid-thickness surface and displayed on an inflated HCP surface (https://balsa.wustl.edu/reference/show/pkXDZ).) **(c)** Pearson correlation (see text for highly similar Spearman values) between MD regions for the H>E contrast, computed across individuals (n = 216). **(d)** Stability of MD H>E effect sizes across runs across individuals (n = 216). **(e)** MD H>E effect sizes and behavior relationship: larger MD H>E effect sizes are associated with better accuracy (*left*) and faster RTs (*middle*) in the spatial WM task (n = 216), as well as higher IQ scores (n = 114) (*right*) as measured by an independent test (KBIT).

To estimate the fROIs’ responses to the Hard and Easy conditions, we used an across-run cross-validation procedure (Nieto-Castañón & Fedorenko, 2012) to ensure that the data used to identify the fROIs are independent from the data used to estimate their response magnitudes (Kriegeskorte, Simmons, Bellgowan, & Baker, 2009). To do this, the first run was used to define the fROIs and the second run to estimate the responses. This procedure was then repeated using the second run to define the fROIs and the first run to estimate the responses. Finally, the responses were averaged across the left-out runs to derive a single response magnitude estimate for each participant in each fROI for each condition. Finally, these estimates were averaged across the 18 fROIs of the MD network to derive one value per condition for each participant (see **Fig. 2c** for evidence of strong inter-region correlations in effect sizes, replicating Mineroff et al., 2018). (An alternative approach could have been to examine fROI *volumes* – the number of MD-responsive voxels at a fixed significance threshold – instead of effect sizes. However, first, effect sizes and region volumes are strongly correlated; and second, effect sizes tend to be more stable within participants than region volumes (Mahowald & Fedorenko, 2016)).

### Language fROIs definition and response estimation

To define the language fROIs, we used a set of six functional masks that were generated based on a group-level representation of data for the Sentences>Nonwords contrast from a large set (n=220) of participants (e.g., Paunov et al., 2019). These masks included three regions in the left frontal cortex: two located in the inferior frontal gyrus, and one located in the middle frontal gyrus; and three regions in the left temporal and parietal cortices spanning the entire extent of the lateral temporal lobe and going posteriorly to the angular gyrus. Within each masks, we selected the top 10% of most responsive voxels in each individual participant based on the *t-*values for the Sentences>Nonwords contrast. To estimate the fROIs’ responses to the Sentences and Nonwords conditions, we used the across-run cross-validation procedure described above.

### Data availability

Stimuli presentation codes, analysis codes and data (behavioral measures, activation beta estimates and brain maps) are available at https://osf.io/2tw6j/. Access to raw data can be requested by e-mailing E.F.

## Results

### Reliability of behavioral measures

Behavioral performance on the spatial WM task was as expected: individuals were more accurate and faster on the easy trials (accuracy=92.22% ± 7.88%; RT=1.20s ± 0.23s) than the hard trials (accuracy=77.47% ± 11.10%, *t*(215)=-23.23, *p*<0.0001, Cohen’s *d*=1.53 (effect sizes are based on the two-tailed independent samples *t*-test); RT=1.49s ± 0.25s, *t*(215)=-26.14, *p*<0.0001, Cohen’s *d*=-1.23). Behavioral measures were stable within individuals across runs for overall (averaging across the Hard and Easy conditions) accuracies (*r*=0.66, *p*<0.0001) and RTs (*r*=0.81, *p*<0.0001). In contrast, difference scores (Hard > Easy) were less stable for both accuracies (*r*=0.26, *p*<0.0001) and RTs (*r*=0.46, *p*<0.0001) (**Fig. 1**). To further validate overall scores as a reliable individual measure (i.e., stable across runs within an individual), we tested their correlation with IQ scores, a well-established stable measure, in the subset of subjects (n=114) that performed the IQ KBIT test. Indeed, IQ scores correlated with overall but not difference accuracy scores (*r*(IQ vs. overall)=0.35 vs. r(IQ vs. H>E)=0.0033) whereas the correlations were similar for RTs (*r*(IQ vs. overall)=-0.21 vs. *r*(IQ vs. H>E)=0.22). Thus, in the critical brain-behavior analyses below, we used overall accuracies and RTs rather than the H>E measures, because the former are more stable within individuals as demonstrated by their high correlation across runs and correlation with the well-established stable IQ measure. Furthermore, the H>E behavioral measures might contain a non-linearity, such that smaller between-condition differences are observed in both high performers (when performance is close to ceiling) and low performers (when performance is close to chance).

### MD network activity and behavior

As expected (Fedorenko et al., 2013), each of the eighteen MD fROIs individually, as well the MD network as a whole (averaging across fROIs), showed a highly robust H>E effect across participants separately in each run (*t*s(215)>11.54, *p*s<0.0001, Cohen’s *d*=0.79-1.54). Individual differences in the MD H>E effect sizes were also stable across runs for each MD fROI individually (*r*s=0.60–0.80) and when averaging across fROIs (*r*=0.74, *p*<0.0001; **Fig. 2d**). We used the H>E contrast as it was more stable than task>fixation contrasts (H>fix *r*=0.65 and E>fix *r*=0.31). This greater stability of the H>E contrast plausibly reflects the fact that it factors out variability due to state differences, thus honing in on the relevant variability, related to the level of the MD network’s activity. For each participant, we averaged the H>E effect size across the 18 MD fROIs to derive a single measure because the H>E effect sizes were strongly correlated across the 18 regions (*r*s*=*0.45-0.88; **Fig. 2c**), replicating Mineroff et al., 2018, and in line with general evidence of the MD brain regions forming a tightly functionally integrated system (Assem et al., 2020; Blank et al., 2014; Paunov et al., 2019).

To ensure that the stability of the MD H>E effect size did not depend on the particular details of the fROI definition (i.e., top 10% of most responsive voxels within the masks), we also extracted the effect sizes from the fROIs defined as the top 20% and top 30% of most responsive voxels. The extracted H>E effect sizes were almost perfectly correlated with those extracted from the top 10% fROIs (20% vs 10%, r=0.99, p<0.0001; 30% vs 10%, r=0.98, p<0.0001). Thus, we proceed to use the H>E effect sizes extracted from the original (10%) fROIs.

For each participant, we used behavioral measures from the spatial WM task (overall accuracies and RTs), and one brain activation measure (H>E effect sizes averaged across the 18 MD ROIs). The critical analyses revealed that larger MD H>E effect sizes were associated with more accurate (Pearson’s *r*=0.44, Spearman’s *r*=0.42, both *p*s<0.0001) and faster (Pearson’s *r*=-0.29, Spearman’s *r*=-0.29, both *p*s<0.0001; **Fig. 2e**) performance. To further test the predictive power of MD H>E effect sizes, we cross-compared brain-behavior relationships across runs (Dubois & Adolphs, 2016) and found that MD H>E effect sizes in run 1 correlated with both accuracies (Pearson’s *r*=0.34, Spearman’s *r*=0.33, both *p*s<0.0001) and RTs (Pearson’s *r*=-0.22, Spearman’s *r*=-0.26, both *p*s<0.0001) in run 2, and MD H>E effect sizes in run 2 correlated with accuracies (Pearson’s *r*=0.40, Spearman’s *r*=0.38, both *p*s<0.0001) and RTs (Pearson’s *r*=-0.27, Spearman’s *r*=-0.27, both *p*s<0.0001) in Run 1.

Next, to test the generalizability of the relationship between MD activation and behavior, we asked whether MD H>E effect sizes explain variance in fluid intelligence, as measured with the Kaufman Brief Intelligence Test (KBIT) (Kaufman & Kaufman, 2014) in a subset of participants (n=114). Indeed, larger MD H>E effect sizes were associated with higher intelligence quotient (IQ) scores (Pearson’s *r*=0.34, *p*<0.0002, normalized R^2^(R^2^_H>E vs IQ_ /R^2^_H>E reliability_)=21%; Spearman’s *r*=0.41, p<0.0001 **Fig. 2e**). This relationship was still significant after controlling for WM accuracy using a partial correlation analysis (Pearson’s *r*=0.26, *p*=0.0061; Spearman’s *r*=0.34, p=0.0003), suggesting that MD activity explains unique variance captured by the fluid intelligence test over and above any shared working memory component between the test and the task.

These results thus support a positive association between MD activity and fluid cognitive abilities. In the next section we assess the selectivity of this MD-behavior relationship.

### Language network activity and behavior

Does the strength of brain activity outside of the MD network explain variance in executive abilities? We tested the selectivity of the MD-behavior relationship by examining another large-scale network implicated in high-level cognition: the fronto-temporal language-selective network in the left hemisphere (Fedorenko et al., 2011).

We extracted the language network’s activity during a reading task (Fedorenko et al., 2010) (Sentences>Nonwords (S>N) contrast; **Fig. 3a**). Similar to MD H>E effect sizes, language S>N effect sizes were highly stable across runs for each language fROI individually and averaging across fROIs (*r*=0.83, *p*<0.0001; **Fig. 3b**), in line with prior work (Mahowald & Fedorenko, 2016; Mineroff et al., 2018).

**Figure 3.**
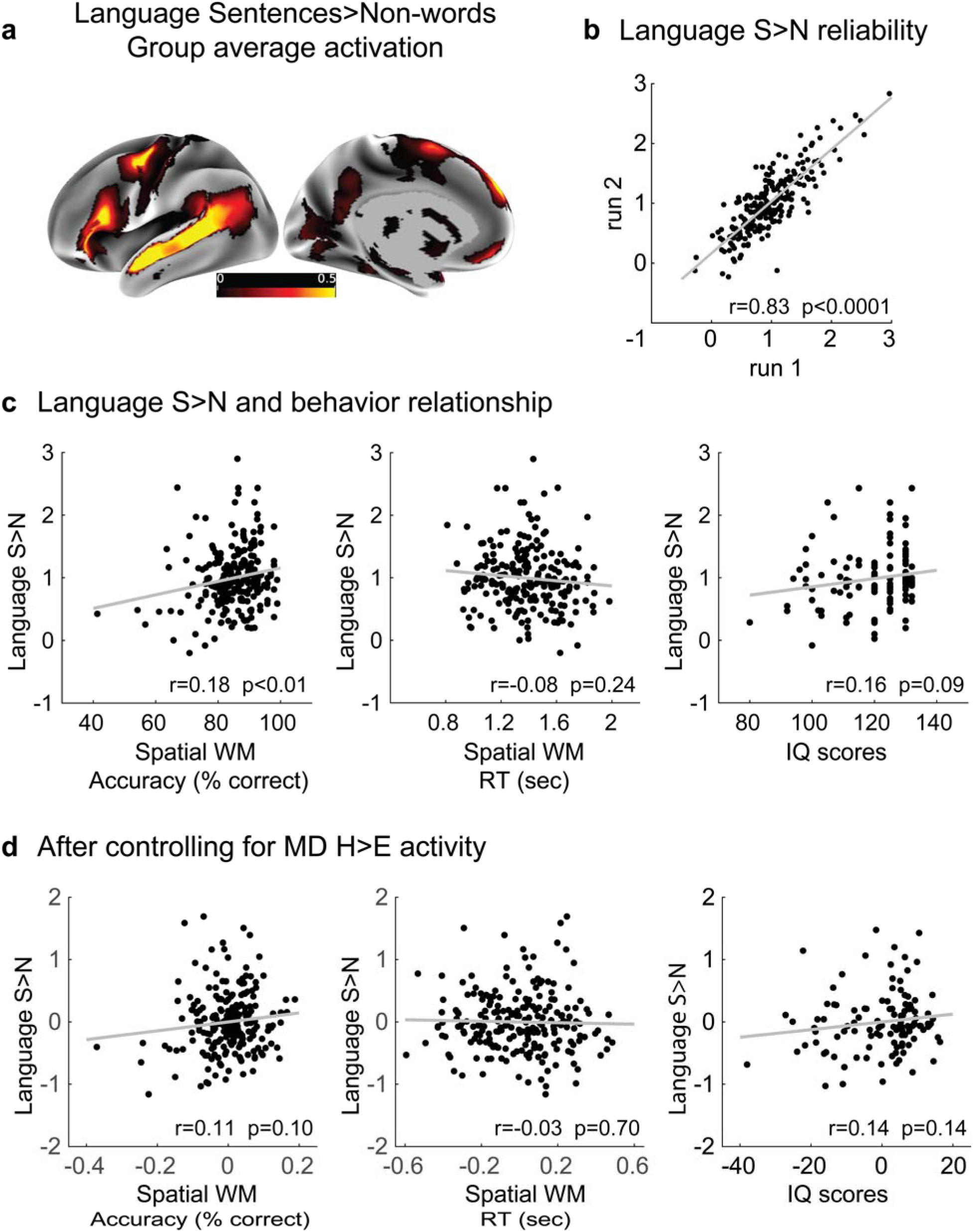
Language network activity and behavior. **(a)** Surface projection of the volumetric unthresholded group average activation map (beta estimates) for the language Sentences>Nonwords (S>N) contrast. **(b)** Stability of language S>N effect sizes across runs across individuals (n=216) (Pearson correlations are used in the figures; see text for highly similar Spearman values). **(c)** Language S>N effect sizes and behavior relationship: larger language S>N effect sizes are weakly associated with better accuracy in the spatial WM task (*left*) and higher IQ scores (*right*), but not RTs in the WM task (*middle*). **(d)** Language S>N effect sizes and behavior relationship, controlling for MD H>E effect sizes: the weak relationships between language S>N effect sizes and behavior observed in (c) are now abolished.

Larger language S>N effect sizes were weakly associated with more accurate (Pearson’s *r*=0.18, *p*<0.01; Spearman’s *r*=0.17, *p*=0.01) but not faster (Pearson’s *r*=-0.08, p=0.24; Spearman’s *r*=-0.10, *p*=0.14) performance on the spatial WM task **(Fig. 3c**). We also observed a weak trend for a relationship between S>N effect sizes and IQ scores (Pearson’s *r*=0.16, *p*=0.09; Spearman’s *r*=0.15, *p*=0.11) (**Fig. 3c**). Critically, however, controlling for the size of the MD H>E effects, in a partial correlation analysis, abolished the associations between language S>N effect sizes and the behavioral measures (spatial WM accuracies: Pearson’s *r*=0.11, *p*=0.10, Spearman’s *r*=0.18, *p*=0.09; IQ scores: Pearson’s *r*=0.14, *p*=0.14, Spearman’s *r*=0.11, *p*=0.25; **Fig. 3d**). In contrast, controlling for the size of the language S>N effects did not affect the relationship between MD H>E effect sizes and the behavioral measures (spatial WM accuracies: Pearson’s *r*=0.42 cf. *r*=0.44; spatial WM RTs: Pearson’s *r*=-0.27 cf. *r*=-0.29; IQ scores: Pearson’s *r*=0.34 cf. *r*=0.35; all *p*s<0.001).

In line with findings from brain lesion studies, these results confirm the selective relationship between the MD network and executive functions / fluid intelligence.

### Effect of sample size and reliability of the fMRI activity on brain-behavior associations

In a further attempt to explain discrepancies in the prior literature (e.g., some studies finding that stronger MD activity is associated with better executive abilities, but other studies finding the opposite pattern, as discussed in the Introduction), we examined the effects of sample size and reliability of the fMRI effect sizes on the brain-behavior relationships (Gelman & Carlin, 2014). We used two indices of MD activity that differed in their reliability – (1) MD H>E effect size used in the main analysis above (a highly reliable measure, with the across-runs correlation of Pearson’s r=0.74) and (2) MD E>Fix effect size (a less reliable measure, with the across-runs correlation of Pearson’s r=0.31) – and examined their relationship to IQ scores.

Samples of different sizes (ranging from 10 to 110, in increments of 10) were randomly selected from our set of 114 participants. For each sample, we computed a correlation between each of the two activity measures and IQ scores. This process was repeated 1,000 times per sample size. The resulting correlations were then examined for their sign, size, and significance. The results, shown in **Fig. 4 (left)**, clearly demonstrate that a combination of small samples and brain activity measures of low reliability (e.g., MD E>fix effect size), like those used in many earlier studies, can produce a significant (p<0.05) correlation of the opposite sign to that observed in a larger population (red dots with a negative correlation). This problem is diminished, but not eliminated, when a reliable index like the MD H>E effect size is used (**Fig. 4, right**). The results also demonstrate that inflated correlations that are often observed in small samples are not eliminated even when a reliable activity measure is used.

**Figure 4.**
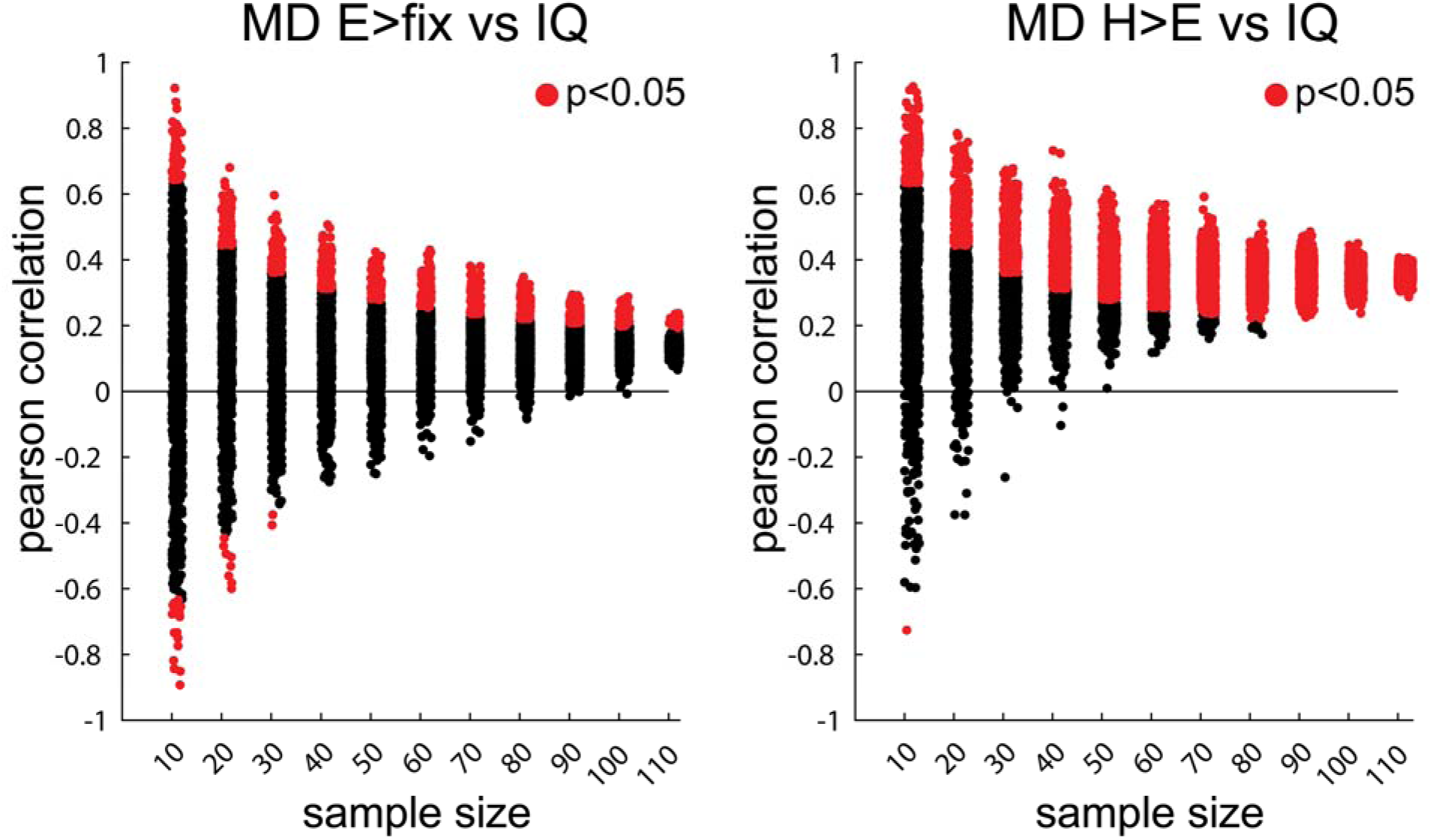
Effects of sample size and the reliability of the brain measure on brain-behavior relationships. On the x-axis in both panels, we show correlations (1,000 per sample) obtained for samples of different sizes. In the left panel, we use a brain activity measure of low reliability (MD E>Fix effect size), and in the right panel, we use a highly reliable brain activity measure (MD H>E effect size). Correlations significant at the p<0.05 level are marked in red.

The results from this analysis also challenge the claim of a negative association between MD activity and performance observed in easier tasks. As demonstrated above, at least in this paradigm, brain activity during a relatively easy executive task was not reliable within individuals across runs. This low reliability could yield correlations of opposite sign. However, even with large sample sizes, the MD E>fix effect size shows a weak positive, not negative, association with IQ scores (**Fig. 4, left**).

## Discussion

In a large set of participants, we examined the relationship between activity in the fronto-parietal “multiple-demand (MD)” network (Duncan, 2010, 2013), on the one hand, and executive abilities and fluid intelligence, on the other. The brain regions of interest were defined in individuals using a functional localizer task (e.g. Fedorenko et al., 2013). We observed a robust positive association between the strength of activation in the MD network and performance on a spatial working memory (WM) task in the scanner, as well as IQ measured independently. We also examined the specificity of this relationship by considering another network important for high-level cognition – the fronto-temporal language-selective network (Fedorenko et al., 2011). Although the strength of activation in this network showed a weak positive association with some of the behavioral measures, these relationships were eliminated once the level of the MD network’s activity was taken into account (controlling for the level of the language network’s activity did not affect the MD-behavior relationships). Finally, we showed how small sample sizes and/or the use of brain activity measures of low reliability, as used in many earlier studies (Dunst et al., 2014; Haier et al., 1988; Lipp et al., 2012; Rypma et al., 2006), could produce inflated and/or the opposite-sign correlations between brain and behavior. To our knowledge, our relatively large sample size, coupled with the participant-specific functional localization approach to defining the regions of interest (Nieto-Castañón & Fedorenko, 2012; Saxe et al., 2006), provides the strongest evidence to date for a positive association between the MD network’s activity and behavioral measures of executive abilities and fluid intelligence. This evidence aligns well with findings from lesion studies that have also reported a selective relationship between fronto-parietal regions and fluid cognitive abilities (Duncan et al., 1995; Glascher et al., 2010; Roca et al., 2010; Warren et al., 2014; Woolgar et al., 2018, 2010).

### Constraints on generality

Some limitations for our study are worth noting (Simons, Shoda, & Lindsay, 2017). First, some researchers have previously tried to explain the discrepancies in the MD-behavior literature by alluding to differences in the age of participants across studies (Reuter-Lorenz et al., 2000; Rypma & Esposito, 2000), arguing that the MD-behavior relationship may change across the lifespan. These changes may be driven by processes like cognitive reserve and brain maintenance in old age (Nyberg & Pudas, 2019; Sala-Llonch, Bartres-Faz, & Junque, 2015; Stern, 2017) or reorganization of neurocognitive architecture in adolescents (Simpson-Kent et al., 2020). The age range in our sample (25^th^-75^th^ percentile = 20-25) is too narrow to evaluate this hypothesis rigorously. That said, the early studies that had motivated this hypothesis a) used small sample sizes (e.g. Rypma and Esposito, 2000), b) used task>fixation activation measures that are likely to be unreliable, and c) did not take into account inter-individual variability in the locations of the MD regions, which may be especially important given the increased variability in the functional architecture of older adults (Geerligs, Tsvetanov, Cam-CAN, & Henson, 2017).

Second, as briefly mentioned in the introduction, some researchers have argued that negative MD-behavior associations can be observed during some easy tasks. For example, a recent study using the HCP n-back task (Barch et al., 2013) demonstrates that whereas MD activations during the 2-back condition are positively associated with general intelligence, MD activations during the 0-back condition show a negative association (Sripada et al., 2020). It is plausible that our easy condition is more cognitively demanding than the 0-back condition, and that is why we did not observe a negative correlation between the E>Fix activations and IQ scores (**Fig. 4, left**).

More broadly, there are situations where improvement in performance is associated with decreases in MD activity e.g. in paradigms with extended motor skills practice (Dayan & Cohen, 2011; Steele & Penhune, 2010) or task rules practice (Hampshire et al., 2019; Ruge & Wolfensteller, 2010). In such cases, efficient performance is plausibly mediated by re-configuration of brain processes. Extended practice can be conceived as a shift from a novel (hard) task to a routine (easy) task. Shifts from hard to easy tasks are known to be associated with anterior to posterior shifts in peak MD activations (Assem et al., 2020; Badre, 2008; Crittenden & Duncan, 2014; Shashidhara, Mitchell, Erez, & Duncan, 2019). Thus, MD activation decreases with practice could reflect these hard to easy topographical activation shifts.

Third, our study used MD activity estimates during a single task. An estimate derived from multiple MD tasks may more accurately capture the variability in the MD network’s engagement across individuals. Similarly, our measure of fluid intelligence was derived from a single IQ test (KBIT; Kaufman and Kaufman, 2013). A measure of fluid intelligence based on a diverse battery of executive function tasks may be more reliable. Nevertheless, we note that in our study (a) the size of the correlation we observed (r=∼0.35) is within the range of correlations reported in recent studies that have used a multi-task-based estimate of fluid intelligence (Dubois et al., 2018; Sripada et al., 2020), (b) the relation between MD-IQ survived after controlling for the correlation between IQ and WM performance, highlighting the unique behavioral variance captured by the KBIT test over and above the WM task.

### MD system activation and intelligence

We estimated MD activity using a blocked design experiment, thus averaging across multiple cognitive processes (in our case, encoding of information into working memory, maintaining and dynamically updating it, and finally, retrieving it from working memory at the decision-making step). Temporally finer-grained MD activity estimates at particular steps in an executive-function task may more precisely target the core neural computations that relate to executive abilities / fluid intelligence. For instance, a recent event-related study demonstrated robust MD activity at each of the stages above (Soreq, Leech, & Hampshire, 2019). Stronger MD activation during more difficult tasks is thought to reflect the increased demand on integrating more and/or different kinds of information in a focused attentional structure to solve the task at hand (Assem et al., 2020; Duncan, 2013). For example, in a recent event-related study, individuals with lower intelligence scores, compared to those with higher scores, showed weaker MD activity during the critical step of target detection suggesting a failure to correctly integrate task rules in the attentional structure guiding successful behavior (Tschentscher et al., 2017). Thus, stronger MD activity across an entire block could plausibly reflect less frequent lapses of “attentional focus” – needed for the correct binding of information to solve the task at hand – and thus better behavioral performance.

A general challenge with individual-level estimates from event-related designs is that they are likely to be more noisy / less reliable, although with sufficient data per participant, this limitation could be overcome. An early study (Gray et al., 2003) with 60 participants found a significant difference between higher and lower IQ individuals in MD activity when it was estimated from individual lure trials (in a n-back task) but not when MD activity was estimated across an entire block of trials. In our study, we demonstrate that MD activity estimated from a block of trials carries meaningful variance about individual differences in executive performance and fluid intelligence.

### Relationship of executive abilities with language and other non-MD regions

Studies of brain lesions have demonstrated repeatedly that there is no relation between lesions in the language network and executive abilities (Fedorenko and Varley, 2016; Woolgar et al., 2018; cf. Baldo et al., 2010). To our knowledge, this is the first study to investigate the relationship between brain activity in the language network and executive abilities / intelligence employing a large sample size and individual-subject fROIs. In line with lesion findings, we show that controlling for MD activity abolishes any relationship between activity in the language network and spatial WM performance and IQ scores. The weak language-behavior association observed prior to controlling for MD activity is plausibly related to a trait factor like vascularization, or a state factor like arousal.

More generally, as we have briefly alluded to in the introduction, several studies have linked executive abilities and fluid intelligence to diverse structural and functional brain measures, including outside the boundaries of the MD network. For example, a recent large-scale study using the UK Biobank dataset (n=∼30,000) reported that total brain volume, as well as multiple global measures of grey and white matter macro- and microstructure (especially, in older participants), explained substantial variance in fluid intelligence (Cox, Ritchie, Fawns-Ritchie, Tucker-Drob, & Deary, 2019). Another study used the HCP task fMRI dataset to show that task-related activations in many brain regions correlates to some extent with general intelligence. However, executive tasks engaging MD regions were the best predictors of individual differences in general intelligence (Sripada et al., 2020), in line with our findings. The relationship among the different neural measures that have been shown to predict variation in fluid intelligence, including the one used in the current study (i.e., the relative increase in the MD activity for a more difficult compared to an easier version of an executive task), is not known. Further studies that assess the reliability of those diverse brain measures, extracted with analysis pipelines that respect inter-individual variability in structure (Kharabian Masouleh, Eickhoff, Hoffstaedter, & Genon, 2019) and function (Coalson, Essen, & Glasser, 2018; Nieto-Castañón & Fedorenko, 2012), and direct comparisons among those measures can help clarify their unique and shared contributions to explaining variability in executive abilities and intelligence. Given the complexity of human reasoning abilities, multiple brain processes likely contribute, but we suggest that the MD network is a key player governing individual differences in fluid intelligence and executive abilities, in line with the fact that damage to MD structures selectively and robustly predicts intelligence losses.

### Implications for future studies

There are many long-recognized challenges facing brain-behavior individual-differences studies (Dubois & Adolphs, 2016). In the introduction we highlighted the critical role of individually defined functional regions to correctly delineate brain-behavior relationships. Another challenge concerns small sample sizes. Our results demonstrate that typical sample sizes (n=10-30) in neuroimaging studies can produce misleading and highly inflated brain-behavior correlations. This presents a significant challenge for laboratory-based research, clinical studies with difficult to recruit patients and longitudinal studies that opt for multiple scanning sessions at the expense of increasing sample size.

We also demonstrate how unreliable brain or behavioral measures (i.e. not stable within an individual across runs/sessions) can result in invalid and inflated correlations. Reliability can also be compromised by using tasks that do not generate enough between-individuals variance (Hedge, Powell, & Sumner, 2018). This is a general challenge facing integrating experimental and individual differences approaches. For example, response inhibition tasks (e.g. stroop, Go/No-Go) produce replicable experimental effects yet studies on individual differences in performance on these tasks commonly fail to group them in a single construct (Hedge et al., 2018; Rey-Mermet, Gade, Souza, von Bastian, & Oberauer, 2019) or relate them reliably to common brain mechanisms (Rosenberg et al., 2019; Wager et al., 2005).

### Conclusions

Against a backdrop of contradictory prior findings, we demonstrate a robust positive and selective association between the MD network’s activity level, on the one hand, and executive abilities and fluid intelligence, on the other. Our analyses also help resolve some of the prior contradictions in the literature. Given its high reliability, the MD activity measure used here, and measures obtained from similarly strong manipulations of cognitive demand, can be used as a neural marker to further probe variability in executive abilities both in the typical population and among individuals with cognitive and psychiatric disorders. This marker can also serve as a promising neural bridge (Braver, Cole, & Yarkoni, 2010) between behavioral variability and genetic variability associated with differences in fluid intelligence (Deary, Spinath, & Bates, 2006; Plomin & Spinath, 2004).

## Acknowledgments

We thank John Duncan, Alex Woolgar, Tamer Demiralp, Aysecan Boduroglu, Burak Guclu, and EvLab members for providing helpful comments on this work. We thank Zuzanna Balewski for setting up the working memory experiment, Matt Siegelman for help with organizing behavioral and neural data, and Hannah Small for help with organizing the demographic data. The authors would also like to acknowledge the Athinoula A. Martinos Imaging Center at the McGovern Institute for Brain Research at MIT, and the support team (Steven Shannon and Atsushi Takahashi). E.F. was supported by NIH awards R00-HD-057522, R01-DC-016607, and R01-DC-016950, and by a grant from the Simons Foundation to MIT’s Simons Center for the Social Brain. M.A. was supported by Cambridge Trust-Yousef Jameel Scholarship.

## Author Contributions (CRediT Taxonomy)

**M.A.** Conceptualization; Data curation; Formal analysis; Software; Methodology; Visualization; Validation; Writing - original draft; Writing - review & editing.

**I.A.B.** Data curation; Formal analysis; Software; Methodology; Investigation; Validation Writing - review & editing.

**Z.M.** Data curation; Formal analysis; Software; Investigation; Project administration.

**A.A.** Methodology; Supervision; Writing - review & editing.

**E.F.** Conceptualization; Data curation; Formal analysis; Funding acquisition; Investigation; Methodology; Project administration; Resources; Software; Supervision; Validation; Roles/Writing - original draft; Writing - review & editing.

